# Initiation of aboveground organ primordia depends on combined action of auxin, *ERECTA* family genes, and PINOID

**DOI:** 10.1101/2022.02.24.481843

**Authors:** Daniel DeGennaro, Ricardo Andres Urquidi Camacho, Liang Zhang, Elena D. Shpak

## Abstract

Leaves and flowers are produced by the shoot apical meristem (SAM) at a certain distance from its center, a process that requires the hormone auxin. The amount of auxin and the pattern of its distribution in the initiation zone determine the size and spatial arrangement of organ primordia. Auxin gradients in the SAM are formed by PIN-FORMED (PIN) auxin efflux carriers whose polar localization in the plasma membrane depends on the protein kinase PINOID (PID).

Previous work determined that ERECTA family genes (ERfs) control initiation of leaves. ERfs are plasma membrane receptors that enable cell-to-cell communications by sensing extracellular small proteins from Epidermal Patterning Factor/EPF-like (EPF/EPFL) family. Here, we investigate whether ERfs regulate initiation of organs by altering auxin distribution or signaling. Genetic and pharmacological data suggest that ERfs do not regulate organogenesis through PINs while transcriptomics data show ERfs do not alter primary transcriptional responses to auxin.

Our results indicate that in the absence of ERf signaling, the peripheral zone cells inefficiently initiate leaves in response to auxin signals and that increased accumulation of auxin in the *er erl1 erl2* SAM can partially rescue organ initiation defects. We propose that both auxin and ERfs are essential for leaf initiation, and that they have common downstream targets. Genetic data also indicate that the role of PID in initiation of cotyledons and leaves cannot be attributed solely to regulation of PIN polarity, and PID is likely to have other functions in addition to regulation of auxin distribution.

**Summary statement:** Auxin is unable to promote cotyledon and leaf initiation in the absence of signaling by ERECTA family receptor kinases and the kinase PINOID.

## INTRODUCTION

Leaves and flowers are formed at the periphery of the shoot apical meristem (SAM) in a well-ordered pattern. Across species, the SAM forms lateral organs at varying rates and geometries, giving rise to the wide variety of plant shapes observed in nature. Diverse phyllotactic patterns and leaf shapes enable optimal light absorption that is fine-tuned to specific environmental conditions (Sarlikioti et al., 2011). In Arabidopsis, consecutive leaves and flowers are formed at 137.5° angles to each other. In long days, leaves and flowers is initiated about every 34 and 12 hours, respectively (Cole et al., 2006).

Lateral organs are initiated at a fixed distance from the center of the SAM (Steeves and Sussex, 1989; Lyndon, 1990; Reinhardt et al., 2000). The molecular mechanism by which lateral organ formation is confined to a narrow ring of founder cells surrounding the central zone of pluripotent cells is not fully understood. It has been proposed that lateral organs initiate at the junction of adaxial and abaxial gene expression in an auxin-dependent manner (Przemeck et al., 1996; Caggiano et al., 2017; Heisler and Byrne, 2020). However, other mechanisms cannot be excluded.

The first sign of organ formation is the development of an auxin maximum in the outermost L1 layer of the SAM (Benková et al., 2003; Heisler et al., 2005; Smith et al., 2006). Polar auxin transport to a specific spot in L1 depletes the neighboring areas of auxin and ensures that primordia are properly spaced (Reinhardt et al., 2003). The distribution of auxin in the L1 founder cells not only establishes phyllotaxy, but also influences the size and shape of forming organs, with higher concentrations of auxin inducing formation of wider leaf primordia (Reinhardt et al., 2000). While there are multiple influx and efflux transporters, the directionality of auxin transport depends primarily on polarly localized PIN-FORMED (PIN) efflux transporters. Out of eight PINs, PIN1 plays the leading role in formation of auxin maxima in the L1 layer of the SAM (Vernoux et al., 2000; Reinhardt et al., 2003; Kierzkowski et al., 2013). The positive feedback loop between auxin and PIN1 is essential to the organization of auxin transport. PIN1 is polarly localized in the plasma membrane toward cells with higher auxin concentration (Reinhardt et al., 2003), and its polarity depends on phosphorylation by several classes of protein kinases (Sauer and Kleine-Vehn, 2019). One critical kinase is PINOID (PID), which phosphorylates several Ser residues in the central hydrophilic loop of PINs and promotes their delivery to the cell apical side (Friml et al., 2004; Dhonukshe et al., 2010).

After initial accumulation in the L1 layer, a gradient of auxin appears in the internal tissues underneath the maximum (Benková et al., 2003). This pattern of auxin accumulation coincides with PIN1 expression, which originally led to the hypothesis that PIN1 is responsible for auxin export from L1 into internal tissues (Reinhardt et al., 2003; Smith et al., 2006); however, later experiments suggested that auxin canalization in internal tissues is independent of PIN1 and might be due to diffusion (Kierzkowski et al., 2013; Ravichandran et al., 2020).

In our previous work, we demonstrated that leaf initiation and phyllotaxy are dependent upon a signaling pathway activated by three *ERECTA* family genes (*ERfs*): *ERECTA*, *ERL1*, and *ERL2* (Chen et al., 2013; Kosentka et al., 2019). *ERf* genes encode leucine-rich repeat (LRR) receptor-like kinases (Shpak et al., 2004) that are localized in the plasma membrane, where they sense ligands from the EPF/EPFL (epidermal patterning factor/EPF-like) family (Hara et al., 2007; Shpak, 2013; Lin et al., 2017; Qi et al., 2017). Leaf initiation is significantly reduced in *er erl1 erl2* and *epfl1 epfl2 epfl4 eplf6* mutants, and rosette leaves arise at almost random angles in *er erl1 erl2* (Chen et al., 2013; Kosentka et al., 2019). These changes in leaf initiation coincide with altered auxin accumulation. Analysis of DR5rev:GFP expression indicated that while more auxin accumulates in the L1 layer of *er erl1 erl2*, it does not form well-defined maxima (Chen et al., 2013). Simultaneously, auxin accumulation is severely reduced in the procambial cells of incipient primordia and in vasculature of leaves and the hypocotyl. In *er erl1 erl2,* expression of PIN1 is increased in the L1 layer and decreased in internal tissues (Chen et al., 2013). As PIN1 expression depends on auxin (Paciorek et al., 2005), it was unclear whether the altered PIN1 accumulation was the cause or the consequence of changed accumulation of auxin and decreased leaf initiation.

In this paper, we have further investigated the role of ERfs in the initiation of lateral organs. Our findings indicate that while auxin accumulates at elevated levels in the meristem of *er erl1 erl2*, its ability to induce leaf initiation is reduced. Analysis of genetic interactions suggested that altered auxin transport is not the main cause of reduced leaf initiation in *er erl1 erl2*, and transcriptomic analysis did not detect a substantial role of ERfs in the primary auxin responses. Simultaneously, we observed that ERfs and PID synergistically promote cotyledon and leaf initiation. The quadruple *pid er erl1 erl2* mutant accumulated elevated levels of auxin in the areas where cotyledons and leaves were supposed to initiate, but these organs were not able to form. Based on these data, we propose that efficient cotyledon and leaf initiation depends not only on formation of auxin peaks, but also on ERf signaling and on additional functions of PID outside of its well-known role in regulation of PIN localization.

## RESULTS

### In *er erl1 erl2,* leaves are induced at a higher auxin concertation threshold

The *er erl1 erl2* mutant accumulates high levels of auxin in the L1 layer of the SAM but it forms leaves at a slower rate (Chen et al., 2013). To investigate the role of auxin in leaf initiation in this mutant, auxin accumulation was altered using an inhibitor of auxin biosynthesis and by adding exogenous auxin. L-Kynurenine (Kyn) is a competitive inhibitor of TRYPTOPHAN AMINOTRANSFERASE OF ARABIDOPSIS1/TRYPTOPHAN AMINOTRANSFERASE RELATED (TAA1/TAR) enzymes, which catalyze the first step of auxin biosynthesis (Stepanova et al., 2008; Tao et al., 2008; He et al., 2011). Growth of wild type and *er erl1 erl2* seedlings on media supplemented with Kyn inhibited leaf initiation consistent with the role of auxin in organ initiation (Fig. 1A). We also observed that *er erl1 erl2* is more sensitive to inhibition of auxin biosynthesis than the wild type. After 5 days of growth on media with 10 μM Kyn, the wild type and *er erl1 erl2* form 1.2- and 3.5-times fewer leaves, respectively. On media with 100 μM of Kyn, the wild type and *er erl1 erl2* form 1.4- and 25-times fewer leaves, respectively. These results indicates that, while *er erl1 erl2* accumulates auxin in the L1 layer, leaf initiation in the mutant is more sensitive to changes in auxin biosynthesis that in the wild type.

**Figure 1.**
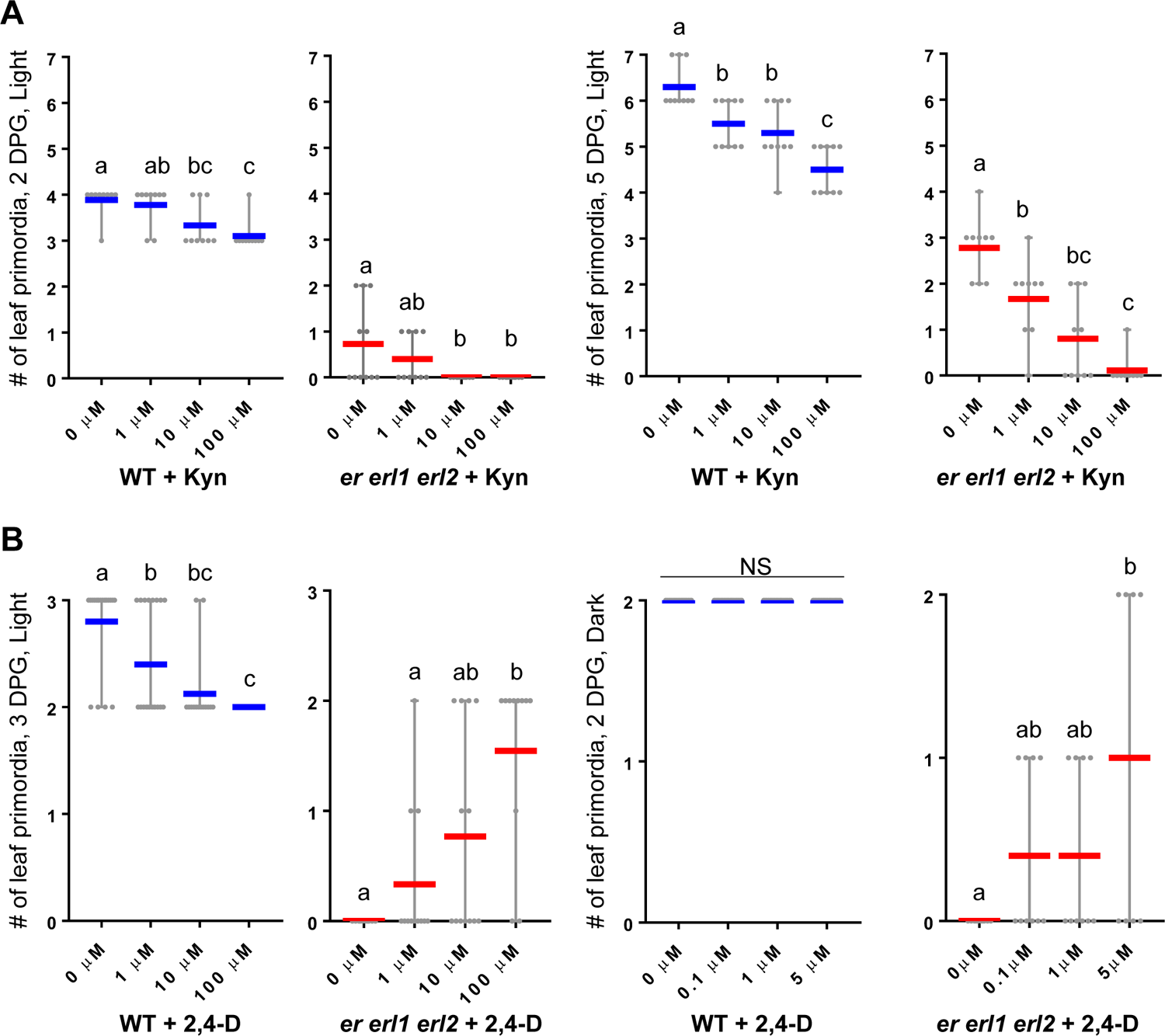
Leaf initiation in the *er erl1 erl2* mutant is limited by auxin availability. **A.** Inhibition of auxin biosynthesis by kynurenine (Kyn) inhibits leaf initiation in both the wild type (WT) and *er erl1 erl2*. **B.** Exogenous auxin (2,4D) promotes leaf initiation in *er erl1 erl2* but not in the WT. **A, B** Seedlings were grown in light or dark as indicated. DPG = days post germination. The mean is indicated as a thick horizontal bar (blue for WT and red for *er erl1 erl2*). The vertical lines designate the maximum and the minimum. Each grey point represents the number of leaves from one individual plant. Different lowercase letters above bars indicate significant difference at P < 0.05, as determined by one-way ANOVA with Tukey’s post hoc test. n=9-20

Next, wild type and *er erl1 erl2* plants were grown in light and in dark on media supplemented with the synthetic auxin 2,4-Dichlorophenoxyacetic acid (2,4-D). In the wild type plants grown in light, the leaf number decreased with increasing concentrations of 2,4-D (Fig. 1B) suggesting that exogenous auxin supplied to the whole seedling can have a negative impact on leaf initiation. In contrast, leaf formation in *er erl1 erl2* was partially rescued by exogenous auxin (Fig. 1B). Etiolated seedlings have a smaller pool of auxin compared to deetiolated (Halliday et al., 2009). In dark, wild type seedlings invariably form two leaves while *er erl1 erl2* is unable to initiate leaves (Fig. 1B). Application of 2,4-D to dark grown seedlings did not alter initiation of leaves in the wild type but *er erl1 erl2* seedlings again showed improved leaf formation. Taken together, these data indicate that compared to the wild type, the *er erl1 erl2* mutant needs higher levels of auxin for efficient leaf initiation.

### A decrease in auxin efflux partially rescues leaf initiation in the *er erl1 erl2* mutant

The auxin efflux transporter PIN1 concentrates auxin into peaks in the SAM and, as a result, plays a key role in leaf and flower formation (Vernoux et al., 2000; Benková et al., 2003; Reinhardt et al., 2003; Heisler et al., 2005). In *er erl1erl2,* PIN1 expression is severely decreased in internal tissues (Chen et al., 2013). To explore whether ERfs regulate leaf initiation through auxin transport, we first investigated genetic interactions between ERfs and PIN1. Both *pin1* and *er erl1 erl2* form fewer leaves than the wild type (Fig. 2A). But when combined, the *pin1* mutation partially rescues leaf initiation defects of the *er erl1 erl2* mutant as determined by two independent experiments (Fig. 2A).

**Figure 2.**
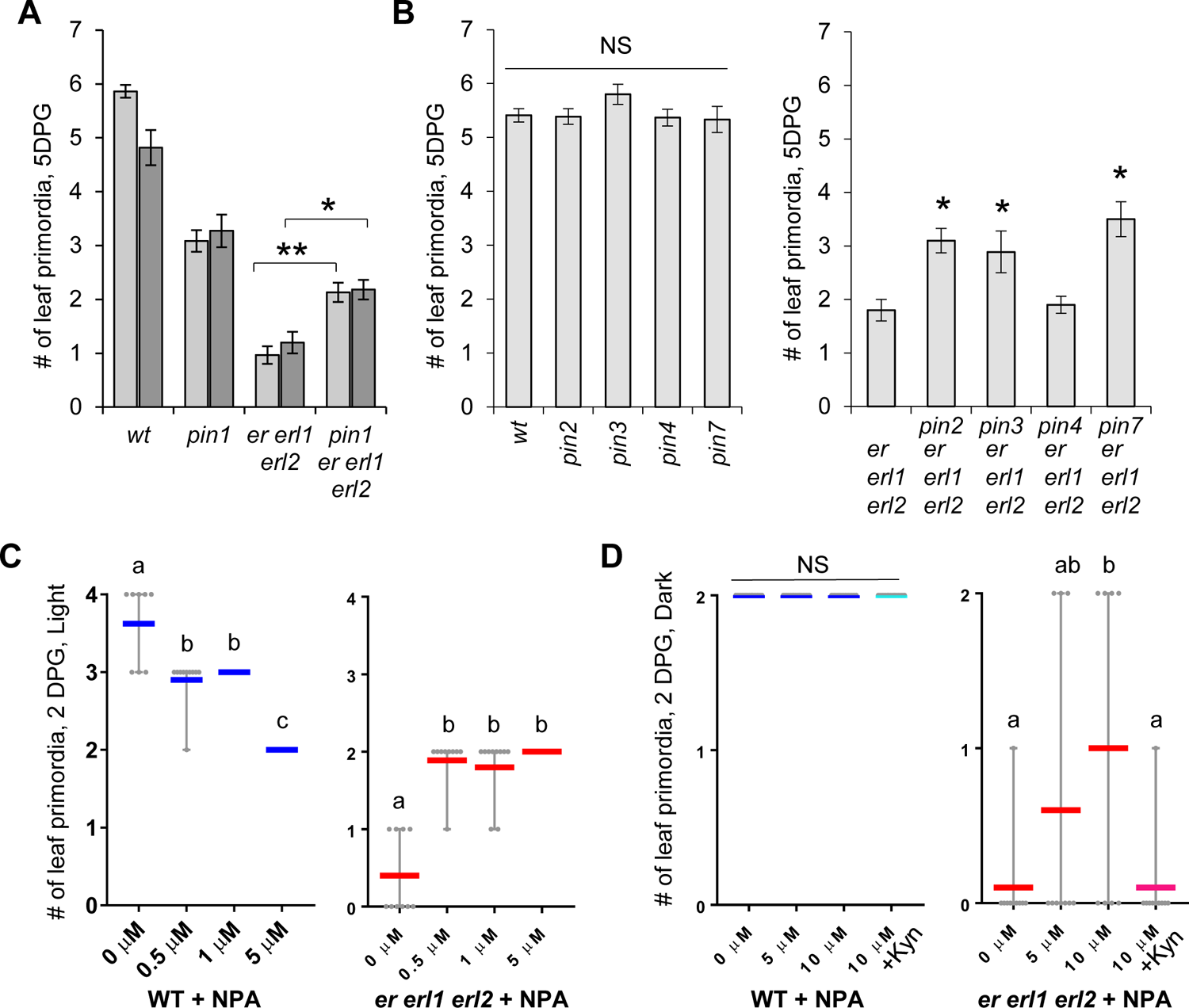
Decrease in auxin transport partially rescues leaf initiation in *er erl1 erl2*. **A.** The *pin1* mutation partially rescues leaf initiation defects of the *er erl1 erl2* mutant. Two independent experiments were performed. Experiment #1 in light grey n=23-30; experiment #2 in dark grey n=10-11; Asterisks above bars indicate significant difference at p<0.005 (*) and p<0.0005 (**) as measured by unpaired two-tailed Student’s t-test. **B.** While the *pin2, pin3, pin4* and *pin7* mutations do not change leaf initiation on their own, *pin2, pin3*, and *pin7* partially rescue leaf initiation of *er erl1 erl2*. n=17-18; An asterisk above a bar indicates significant difference at p<0.01 compared to *er erl1 erl2* as measured by unpaired two-tailed Student’s t-test. NS – no statistical difference compared to the WT. **A** and **B**. Seedlings were grown on solid MS media. Data are mean ± S.E. **C, D.** Inhibition of auxin efflux with NPA reduces leaf initiation in the WT seedlings grown in light and promotes in *er erl1 erl2* in light and in darkness. NPA cannot promote leaf initiation in *er erl1 erl2* seedlings in the presence of 10 mM kynurenine (+Kyn). Seedlings were grown on liquid MS media in light or dark as indicated. DPG = days post germination. The mean is indicated as a thick horizontal bar (blue for WT and red for *er erl1 erl2*). The vertical lines designate the maximum and the minimum. Each grey point represents number of leaves from one individual plant. Different lowercase letters above bars indicate significant difference at p<0.05, as determined by one-way ANOVA with Tukey’s post hoc test. n=8-20.

Out of five PINs implicated in polar auxin transport, only PIN1 is noticeably expressed in the vegetative SAM (Guenot et al., 2012). Leaf initiation is not altered in single *pin2, pin3, pin4* and *pin7* mutants (Guenot et al., 2012) (Fig. 2B), and only *pin4* enhances leaf initiation defects of *pin1* (Guenot et al., 2012). It was proposed that PIN4 contributes to regulation of leaf initiation by altering SAM development during embryogenesis (Guenot et al., 2012). Since higher order *pin* mutations cause severe defects during embryogenesis (Friml et al., 2003), it has not been clearly established whether they play a role in leaf initiation post embryonically. To investigate the relationship between ERf and the other PINs, quadruple *pin er erl1 erl2* mutants were generated. We observed that leaf initiation was partially rescued in *er erl1 erl2* by the addition of the *pin2*, *pin3*, or *pin7* mutations but not by *pin4* (Fig. 2B). These results indirectly implicate *PIN2*, *PIN3*, and *PIN7* in leaf initiation and suggest that reduction of polar auxin transport can partially rescue leaf initiation defects of the *er erl1 erl2* mutant.

To further investigate the role of polar auxin transport in leaf initiation, seedlings were grown for 2 days in light or in darkness on media supplemented with the auxin efflux inhibitor N-1-Naphthylphthalamic Acid (NPA) (Depta et al., 1983; Teale and Palme, 2018). Consistent with previous data (Reinhardt et al., 2000), the wild type formed fewer leaves when exposed to NPA in light (Fig. 2C). In darkness, NPA did not affect the initiation of the first two leaves in the wild type (Fig. 2D). We speculate that the first two leaves are initiated during embryogenesis and postembryonic treatment with NPA does not influence this process. In contrast, treatment with NPA, either in light or in darkness, increased leaf initiation in *er erl1 erl2*. This increase did not occur when *er erl1 erl2* was grown with both NPA and Kyn, indicating that the increased leaf formation in response to NPA is dependent on the availability of auxin. In sum, our data indicate that the decreased leaf initiation in *er erl1 erl2* can be partially rescued by inhibition of auxin efflux.

### Inhibition of auxin efflux leads to increased auxin accumulation in the L1 layer of *er erl1 erl2*

Our previous analysis of DR5rev:GFP expression indicated that 8- to 10-day-old *er erl1 erl2* seedlings had elevated, sheet-like auxin activity in the L1 layer of the SAM and reduced auxin responses in the internal layers (Chen et al., 2013). To investigate auxin accumulation at an earlier developmental stage, DR5rev:GFP expression was examined in 1-day-old seedlings. In the wild type, DR5rev:GFP expression was restricted to narrow regions of the periphery of SAM, a site of incipient leaf primordium, and the expression in L1 was always accompanied by expression in inner tissues (Fig. 3A). In *er erl1 erl2,* we observed an absence of signal in the meristem center and patchy, broad expression in the periphery of the SAM that occasionally correlated with a forming primordium (Fig. 3B and C). The expression was often higher closer to the boundary region and on the abaxial side of primordia (Fig. 3B). The DR5rev:GFP expression was broader than normal at the tips of primordia and was never observed in the inner tissues of *er erl1 erl2*.

**Figure 3.**
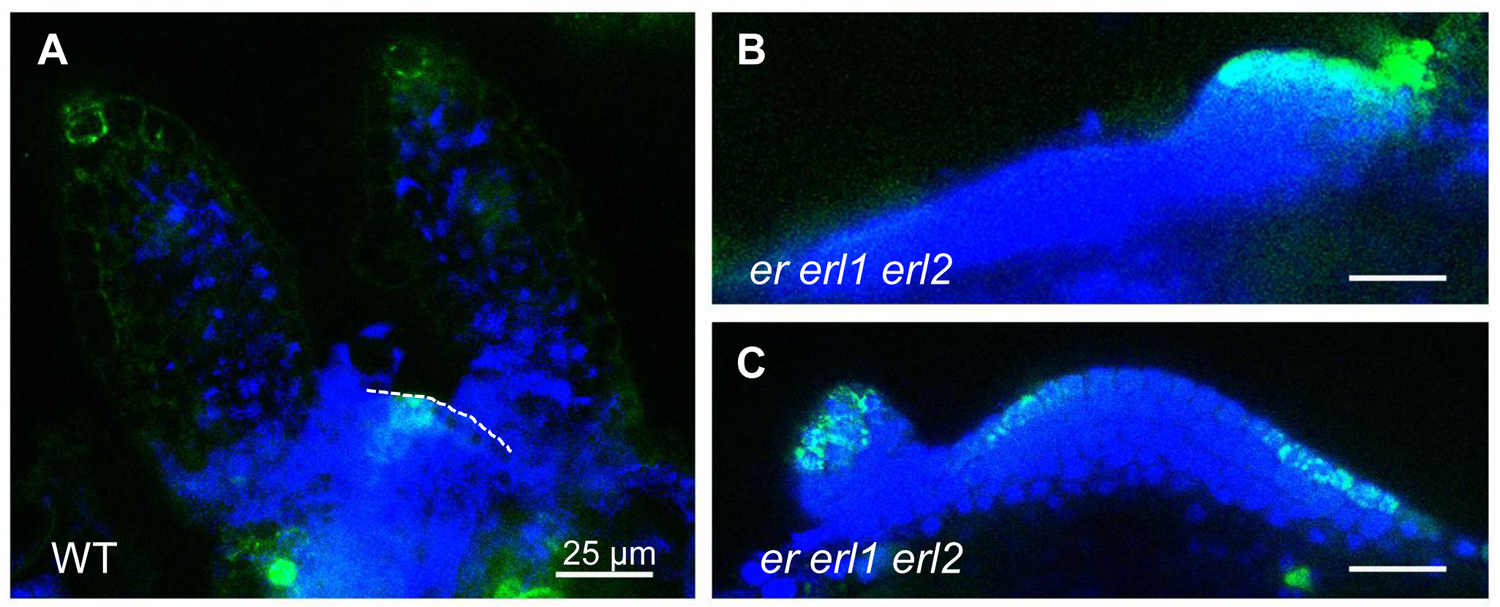
Auxin distribution and sensing is altered in *er erl1 erl2* during early stages of seedling development. Representative confocal images depict auxin localization and perception in the WT and *er erl1 erl2* in 1 DPG seedlings as determined by confocal microscopy using the *DR5rev:GFP* reporter. **A.** In the WT SAM, auxin accumulates narrowly in an incipient leaf primordium in the L1 layer and procambium. The surface of the SAM is indicated by a dashed line. **B-C.** In *er erl1 erl2*, DR5rev:GFP is broadly expressed in the epidermal layer of the SAM but not in the procambium. Its expression is sometimes patchy and often does not coincide with primordia formation. **A-C.** Green = DR5rev:GFP. Blue = DAPI. All images are under the same magnification.

Hypothetically, an absence of auxin in internal tissues could be the cause of decreased leaf initiation. Reduction of auxin efflux promotes leaf initiation in *er erl1 erl2*. Does it alter auxin accumulation in L1, in internal tissues, or in both? To answer this question, changes in DR5rev:GFP expression in response to treatment with 5 μM NPA were examined in one-day-old *er erl1 erl2* seedlings (Fig. 4A-D). The brightness of the GFP signal was measured starting from the boundary of the SAM to its center (Fig. 4C-D). We observed that NPA treatment slightly increased the width of the *er erl1 erl2* SAM (Fig. 4C-D). This is not specific to this mutant as a similar response was previously observed in the wild type (Zhang et al., 2020). Our analysis uncovered an increased auxin accumulation in the L1 layer and no statistically significant change in the L2 layer (Fig. 4C and D). In addition, when the DR5rev:GFP fluorescence was compared in *pin1 er erl1 erl2* and *er erl1 erl2,* we observed an increased signal primarily in the L1 cells of the meristem boundary in the quadruple mutant (Fig. 4A and E). Taken together, our data indicate that inhibition of auxin efflux through genetic or pharmacological methods leads to increased auxin accumulation in the L1 layer of the *er erl1 erl2* mutant.

**Figure 4.**
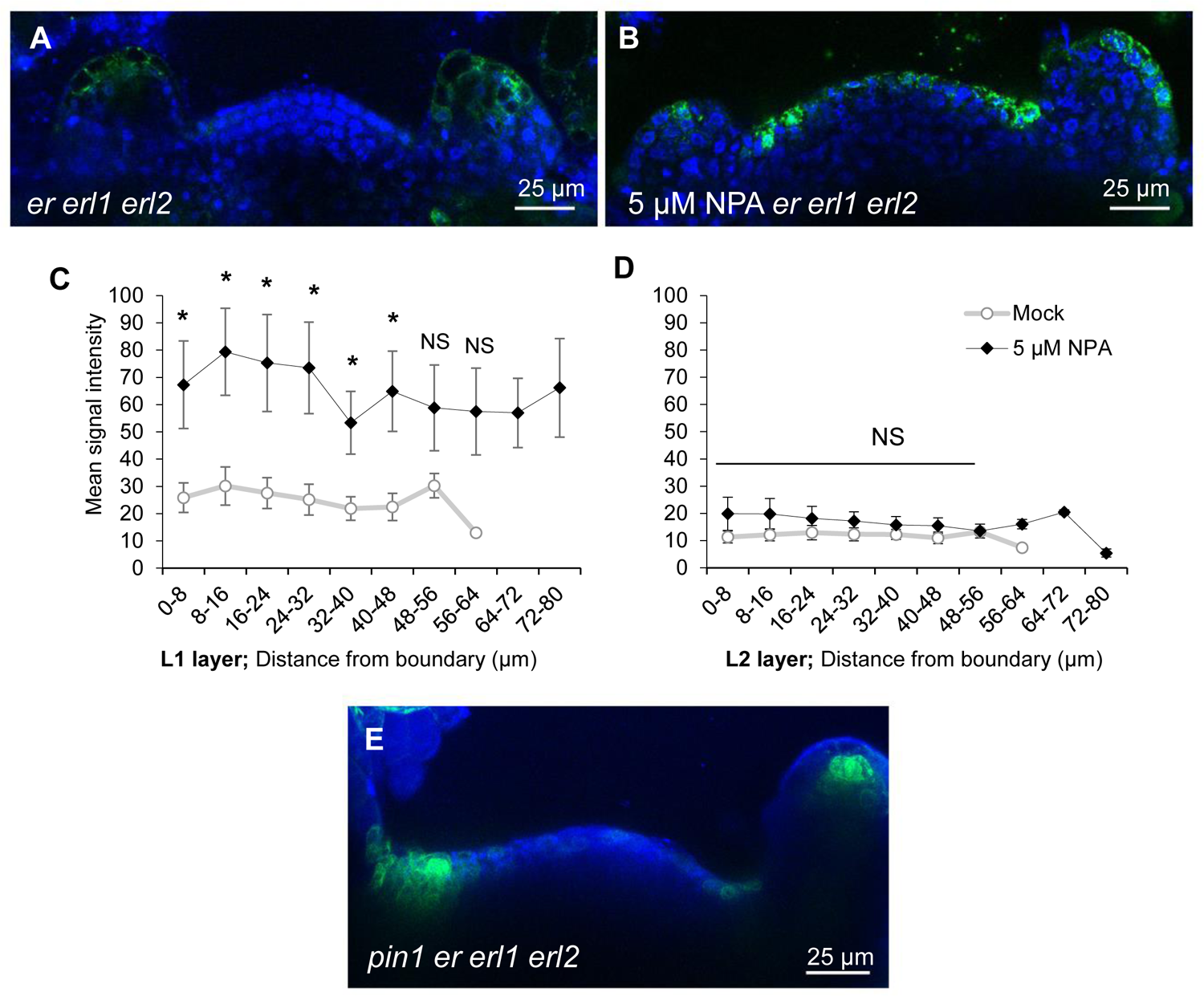
Inhibition of auxin efflux increases auxin accumulation in the L1 layer of *er erl1 erl2*. **A-D.** In response to NPA, auxin accumulates in the L1 layer. **A and B**. Representative confocal images of 1 DPG mock and NPA treated *er erl1 erl2* seedlings expressing DR5rev:GFP. Images are under the same magnification. **C and D.** Quantitative measurements of DR5rev:GFP fluorescence in the L1 (left) and L2 (right) layers of the *er erl1 erl2* SAM. Data points are mean signal intensity ±S.E. Asterisks above data points indicate significant difference at p<0.05 as measured by unpaired two-tailed Student’s t-test. NS = not significant. n=6-14 **E.** A representative confocal image of DR5rev:GFP expression in the *pin1 er erl1 erl2* SAM indicates elevated auxin accumulation in the boundary region of the SAM. 1 DPG.

### In *er erl1 erl2,* auxin can rescue leaf initiation when perceived in either external or internal SAM layers

To test whether increased auxin accumulation in the L1 layer is sufficient to rescue leaf initiation in *er erl1 erl2*, we expressed the modified auxin receptor ccvTIR1 (Uchida et al., 2018) under the L1 specific *ATML1* promoter (Lu et al., 1996). Due to an amino acid substitution in the binding site, the ccvTIR1 receptor cannot sense endogenous auxin; it is activated by synthetic convex IAA (cvxIAA). The presence of a bulky aryl group prevents binding of cvxIAA to endogenous TIR1/AFB receptors. When *ATML1::ccvTIR1* transgenic plants were grown on media containing cvxIAA, leaf initiation was increased in the *er erl1 erl2* background (Fig. 5A) confirming that an increase in auxin signaling in the L1 layer can improve leaf initiation in that mutant. Interestingly, in the wild type or *er erl2* backgrounds we observed a small but statistically significant decrease in the number of leaves in response to cvxIAA (Fig. 5A). We speculate that this might be related to a decrease in the space available for formation of new leaves due to formation of broader leaves, their fusion, and in extreme cases, formation of cup-shaped leaves (Fig. 5D). As expected, a treatment of wild type untransformed plants with cvxIAA did not alter initiation of leaf primordia, confirming that cvxIAA does not regulate leaf initiation through endogenous receptors (Fig. 5C).

**Figure 5.**
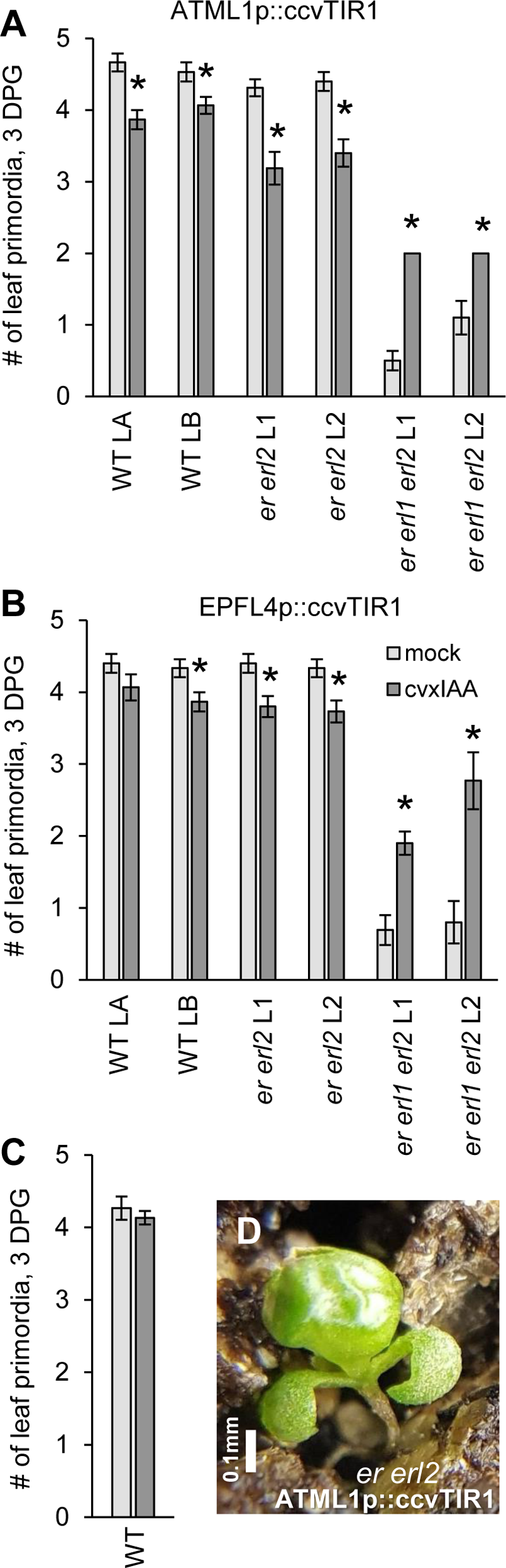
Perception of cvxIAA in the L1 layer or in the internal tissues of the SAM reduces number of leaves in the WT and *er erl2* and increases in *er erl1 erl2*. **A and B.** Comparison of leaf initiation in transgenic plants expressing modified auxin receptor (ccvTIR1) under the epidermal *ATML1* promoter (A) or the SAM internal promoter *EPFL4* (B) in different genotypes as indicated. Seedlings were grown on solid MS media with 50 mM cvxIAA (dark gray bars) or without (light grey bars). Two independent transgenic lines (LA and LB for WT, L1 and L2 for *erf* mutants) were analyzed per genotype. *er erl1* and *er erl1 erl2* of L1 or L2 are siblings. Asterisks above bars indicate significant difference at p<0.05 as measured by unpaired two-tailed Student’s t-test. **C.** cxvIAA does not alter leaf initiation in untransformed WT plants. A-C. Data are mean ± S.E. n=10-15 **D.** Perception of cvxIAA in the L1 layer sometimes leads to formation of a cup-shaped leaf. Three weeks old plant.

To examine the contribution of auxin signaling in the internal layers of the SAM to leaf initiation, the ccvTIR1 receptor was expressed under the *EPFL4* promoter in the L3 layer at the periphery of the SAM (Kosentka et al., 2019). When grown on media containing cvxIAA, leaf initiation was increased in *er erl1 erl2 EPFL4::ccvTIR1* transgenic plants and slightly decreased in the wild type and *er erl2* (Fig. 5B). Overall, our data indicate that upregulation of auxin signaling in either external or internal tissues of the SAM can partially overcome leaf initiation deficiencies of the *er erl1 erl2* mutant and that auxin signaling contributes to leaf initiation not only in the L1 layer but also in the internal layers.

### ERfs and PID synergistically promote organogenesis at the SAM

Polar localization of PIN efflux transporters on the plasma membrane is regulated by the protein kinase PID (Friml et al., 2004; Dhonukshe et al., 2010). Loss of function *pid* mutants greatly resemble *pin1* mutants (Benjamins et al., 2001). To test whether genetic interactions between *ERfs* and *PID* are similar to interactions between *ERfs* and *PIN1,* the quadruple *pid er erl1 erl2* mutant was generated. A strong *pid-3* allele was used in this experiment. Based on the previously described experiments, we expected that *pid*, just like *pin* mutations and NPA, tp partially rescue leaf initiation in the *er erl1 erl2* mutant. However, our predictions were not supported by experimental data.

Wild type plants invariably form two cotyledons (Fig. 6A and B). The e*r erl1 erl2* mutant forms two cotyledons in ∼94% of seedlings, but occasionally initiates one or three cotyledons. Approximately half of *pid* seedlings have an abnormal number of cotyledons: three cotyledons were observed in ∼46% of seedlings and one cotyledon was observed in 3% of seedlings. Unexpectedly, 92% of *pid er erl1 erl2* seedlings did not form any cotyledons (Fig. 6A-C). These data suggest that ERfs and PID are central to cotyledon development, and they function synergistically. The loss of *PIN1* also alters cotyledon initiation: approximately 19% of seedlings in the *pin1-201* allele have an abnormal number of cotyledons (Table S1). Interestingly, the number of cotyledons changes only slightly in the *pin1 er erl1 erl2* mutant compared to *pin1.* We observed an increased number of seedlings with one cotyledon: from 3% in *pin1* to 10% in *pin1 er erl1 erl2* (Table S1). This result suggests a subsidiary role for PIN1 in cotyledon initiation. The *pid er erl1 erl2* mutants not only lacked cotyledons, they also barely formed any leaves and had almost no stem (Fig. 6A and C). The SAM is left exposed in older plants and is easily identifiable by its lack of pigmentation (Fig. 6C). This was an unexpected observation because the *pid-3* mutation on its own does not inhibit leaf initiation or stem elongation (Fig. 6D and E).

**Figure 6.**
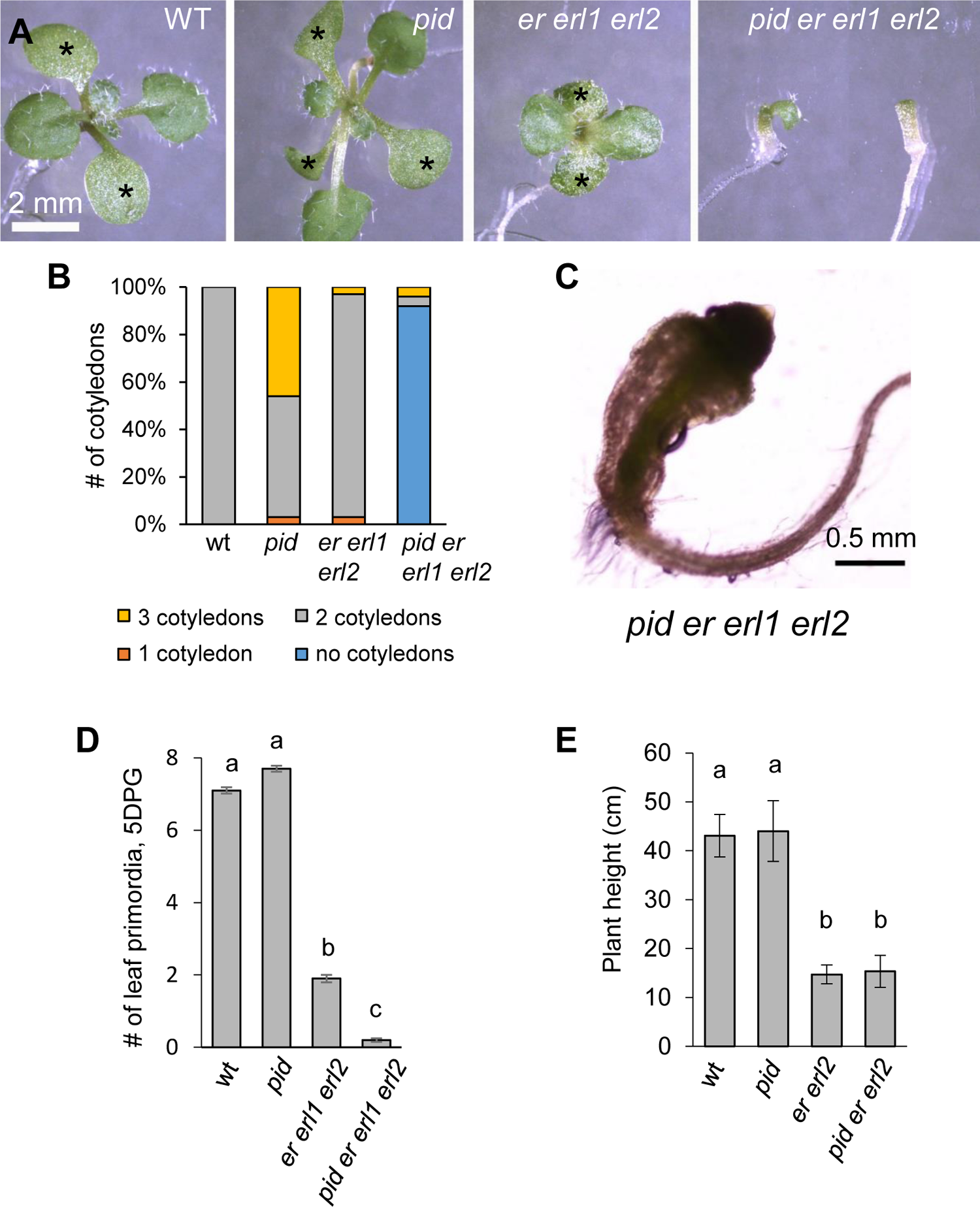
ERfs and *PID* synergistically promote initiation of cotyledons and leaves. **A.** Representative images of 13 DPG seedlings demonstrate a reduced number of cotyledons and leaves in *pid er erl1 erl2*. Cotyledons are indicated by asterisks. All images under the same magnification. **B.** A comparison of cotyledon number in mutants versus WT shows synergistic function of ERf and PID. n=24-442 **C.** A representative image of a one-month-old *pid er erl1 e*rl2 plant demonstrates an absence of cotyledon and leaves and a very short stem. **D.** Number of leaf primordia in 5 DPG seedlings. Data are mean ±S.E; n=10-15. **E.** Average length of the main stem at maturity. Data are mean ±S.D; n=21-37. **D and E**. Different lowercase letters above bars indicate significant difference at P < 0.05, as determined by one-way ANOVA with Tukey’s post hoc test.

Inhibition of polar auxin transport partially rescued leaf initiation in the *er erl1 erl2* background. Why does the *pid* mutation have the opposite effect on initiation of cotyledons and leaves? One possibility is that in the absence of PID auxin efflux transporters move auxin outside of the meristematic area. To test this hypothesis, we analyzed expression of DRrev:GFP during embryogenesis (Fig. 7A). In wild type embryos, most of the signal was observed in the root meristem. There are also maxima at the tips of the cotyledons and a weak signal in the forming vasculature. In *er erl1 erl2*, as was published previously (Chen and Shpak, 2014), the DRrev:GFP signal in the forming vasculature is weaker and the maxima in the cotyledons are either similar or slightly brighter and less focused, while the signal in the root meristem is not altered. A previous analysis of DR5rev:GFP expression in *pid* embryos indicated a weaker and less focused auxin maxima at the tips of developing cotyledons and dramatically reduced auxin accumulation in the developing vasculature (Treml et al., 2005). Our observations are consistent with this previous result (Fig. 7A). In the *pid er erl1 erl2* embryos, we observed very strong expression in the L1 layer in the areas where cotyledons should form and an absence of DR5rev:GFP expression in internal tissues (Fig. 7A). After germination, this mutant maintains an elevated level of auxin in the L1 layer in the periphery of the SAM (Fig. 7B). These data indicate that in the *pid er erl1 erl2* mutant auxin accumulates in the region of cotyledon and leaf formation, but this accumulation is insufficient for organ initiation.

**Figure 7.**
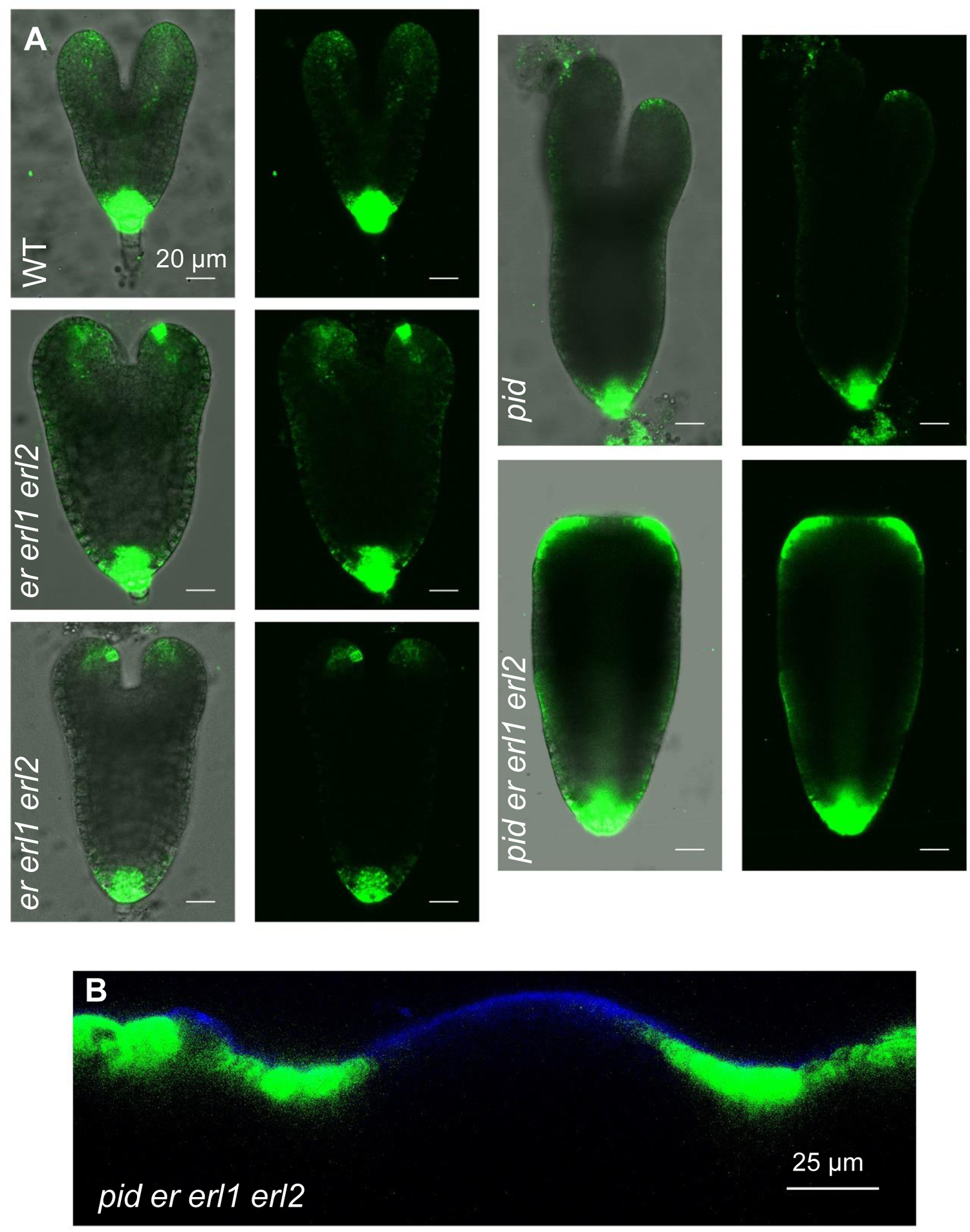
The *pid er erl1 erl2* mutant accumulates elevated levels of auxin in the L1 layer at the periphery of the SAM. **A.** Representative confocal images of DR5rev:GFP expression in WT, *er erl1 erl2*, *pid,* and *pid er erl1 erl2* embryos. All images are under the same magnification. **B.** A representative confocal image of DR5rev:GFP expression in the SAM of 3 DPG *pid er erl1 erl2* seedling. Green = DR5rev:GFP. Blue = DAPI.

We tested whether exogenous auxin could promote organ initiation in *pid er erl1 erl2*. The *pid er erl1 erl2* mutants were grown on media supplemented with 10 μM 2,4-D. After 3 days of exposure to 2,4 D, *pid er erl1 erl2* formed a ring-like tissue outgrowth where the presumptive cotyledons should have been (Fig. 8A). The *pid er erl1 erl2* seedlings left to grow on 2,4-D media for longer than 3 days were severely stunted, possibly due to the inhibitory effect of auxin on root elongation. However, when *pid er erl1 erl2* seedlings treated with 2,4-D for 3 days were moved to standard MS media, after a month we observed a callus-like growth in the apical meristematic region (Fig. 8B). The callus was photosynthetic and formed trichomes and, thus, had some characteristics of leaves. We concluded that in *pid er erl1 erl2* a very high concentration of auxin can induce cell differentiation in the SAM, but proper organs cannot form. Our data indicate that ERfs and PID synergistically promote organogenesis in the periphery of the SAM through a pathway that functions in parallel to auxin.

**Figure 8.**
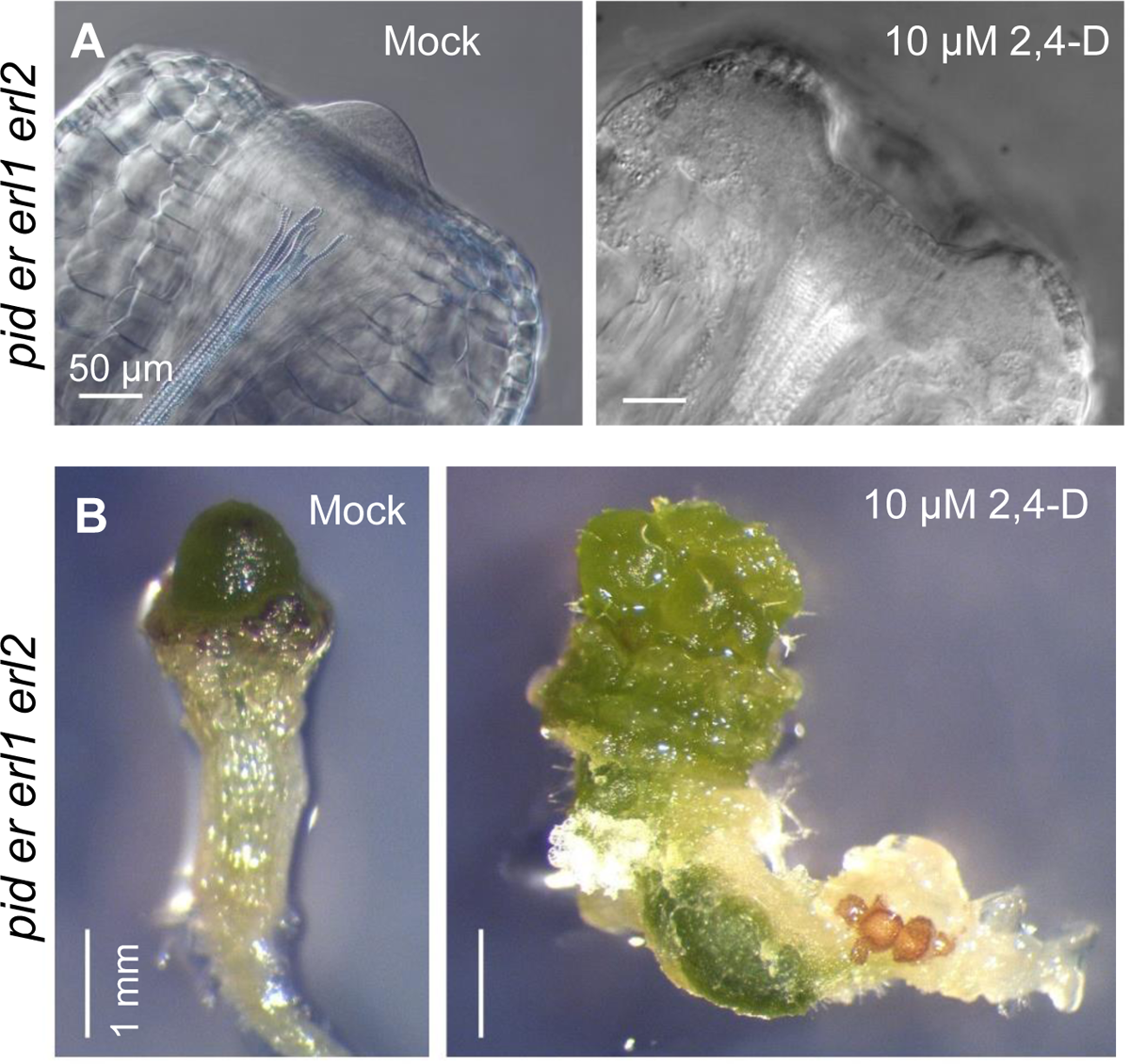
Exogenous auxin induces cell differentiation but not organ formation in *pid er erl1 erl2*. **A.** Representative images of 3 DPG *pid er erl1 erl2* seedlings. Peripheral tissues of the SAM grow and differentiate in response to 2,4-D treatment. **B.** Representative images of 33 DPG *pid er erl1 erl2* plants. When *pid er erl1 erl2* seedlings are transferred to regular media after 16 days on 2,4 D, their meristematic regions form spherical green tissues with trichomes.

### The role of ERfs in organ initiation changes after transition to flowering

Both leaf and flower initiation depend on formation of auxin maxima (Reinhardt et al., 2000). Curiously, mutations in genes regulating auxin transport or sensing have a stronger impact on initiation of flowers compared to leaves. For example, *pin1, pid,* and *mp* form very few flowers, but they initiate rosette leaves at an only slightly reduced rate compared to wild type (Bennett et al., 1995; Przemeck et al., 1996; Gälweiler et al., 1998). On the other hand, the initiation of leaves, but not flowers, is strongly decreased in the *er erl1 erl2* mutant (Chen et al., 2013). To explore the linkage between ERfs and auxin during flower initiation, we investigated genetic interactions between PIN1 and ERfs. Naked, pin-shaped stems form in *pin1* mutants (Fig. 9A). Surprisingly, the loss of *ER* and *ERL2* function partially rescues organ initiation in the *pin1* background. The *pin1 er erl2* mutant forms small, misshapen organs all along the inflorescence stem (Fig. 9B). Organ initiation is rescued further in the *pin1 er erl1 erl2* mutant where randomly positioned carpeloid organs with abundant papillae form at high frequency (Fig. 9C and D).

**Figure 9.**
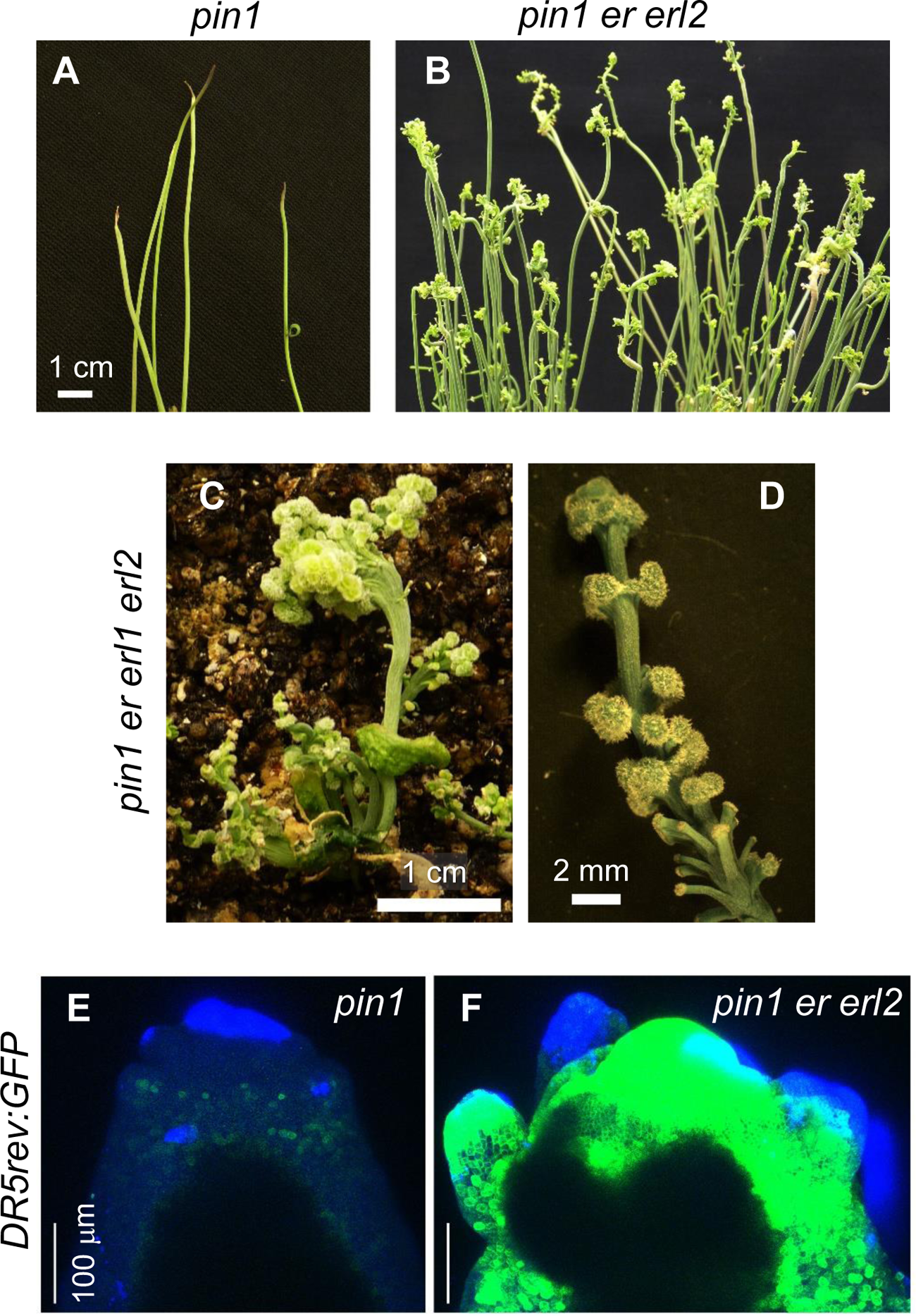
Downregulation of ERf signaling in the *pin1* mutant increases auxin accumulation in epidermis of the inflorescence meristem and partially rescues organ initiation. A and B. Images of inflorescence apexes. Organ initiation is partially rescued in *pin1 er erl2* compared to *pin1*. C and D. The *pin1 er erl1 erl2* mutant is an extreme dwarf which forms numerous randomly positioned carpeloid organs. Plants are 2.5 months old. E and F. Representative confocal images of *DR5rev::GFP* expression in the inflorescence SAM of *pin1* and *pin1 er erl2.* Green = *DR5rev::GFP*. Blue = DAPI.

The inability of *pin1* to form flowers is attributed to the absence of auxin peaks in the inflorescence SAM (Guenot et al., 2012). Our analysis of DR5rev:GFP expression in *pin1* is consistent with that finding; the *pin1* inflorescence SAM had an extremely weak and diffuse fluorescent signal (Fig. 9E). Knockout of *ER* and *ERL2* genes in the *pin1* background dramatically increased auxin accumulation in the meristem; the DR5rev:GFP signal was very high all over the epidermis (Fig. 9F). The loss of *erfs* leads to increased auxin accumulation in the inflorescence SAM which presumably rescues flower initiation due to the *pin1* mutation. In summary, we observed that ERfs have a negative effect on auxin accumulation in the L1 layer of both the vegetative and inflorescence SAM, but they promote formation of lateral organs only before transition to flowering.

The study of *pin1 er erl1 erl2* development also identified a delayed transition to flowering and synergistic regulation of stem elongation by ERfs and PIN1 (Fig. S1). While stem elongation is only slightly reduced in *pin1* compared to the wild type, addition of *pin1* to the *er erl1 erl2* background leads to severe reduction in stem elongation (Fig. S1B and C). Our current and previous work (Chen et al., 2013) suggest that ERfs are necessary for auxin accumulation in internal tissues. As PIN1 promotes auxin transport through the xylem parenchyma and vascular cambium of stem segments (Okada et al., 1991; Bennett et al., 2016), it would be interesting to investigate whether the dramatically decreased elongation of *pin1 er erl1 erl2* internodes is due to auxin deficiency in internal tissues.

### *ERf* genes do not regulate direct auxin responses

The *erf* mutants accumulate a higher amount of auxin in the L1 layer of the SAM, but there is still not enough auxin to efficiently induce formation of leaf primordia. The high expression of DR5rev:GFP in mutants implies that this auxin can be perceived. However, it does not reflect the breadth of auxin responses. Activity of ARFs and the resulting specificity of auxin responses are modulated through cross-talk with other signaling pathways (Chandler, 2016; Lanctot et al., 2020). To investigate whether ERfs modify a subset of auxin responses, we compared gene expression in the wild type and *er erl2* three-day-old seedlings in response to auxin treatment. We chose to investigate the *er erl2* mutant because it forms a reduced number of leaves by five days (Chen et al., 2013), but at three days it is morphologically very similar to the wild type. Thus, any changes in gene expression should not be due to differences in tissue ratios but instead to the disrupted ERf signaling pathway.

Comparison of gene expression between the mutant and the wild type identified an at least 2-fold change in expression of more than 500 genes (Fig. S2A). GO terms for genes upregulated in the mutant showed enrichment in photosynthesis, chloroplast organization, protein targeting to chloroplasts, and starch metabolism. Curiously, while *er erl2* seedlings are not perceptibly different in color, *er erl1 erl2* seedlings are considerably darker green and that characteristic can be used for phenotypic identification in addition to stomata clustering.

Downregulated genes in the mutant showed enrichment for cell wall organization and biogenesis and cell growth (Fig. S2B). Our previous phenotypic analysis connected ER with enhancement of the cell growth rate (Bundy et al., 2012). Consistent with the inhibitory role of ERfs in stomata formation, expressions of many genes linked with this process, such as *TMM*, *EPF1*, *EPF2*, *STOMAGEN*, *SDD1*, *POLAR*, *BASL*, *SCREAM1*, and *FAMA,* were increased in the mutant (Fig. S2C). A thirty-minute treatment with auxin did not alter expression of these genes, suggesting that they are not direct auxin targets. Previous research identified *STOMAGEN* as a direct target of MONOPTEROS (MP) and reported a decrease in *STOMAGEN* expression after 30 minutes of 2,4D treatment (Zhang et al., 2014). But neither our study nor several other examinations of direct auxin targets (Winter et al., 2007; Uchida et al., 2018) support this finding. Consistent with the previous report that ERf genes are upregulated to compensate for loss of family members (Pillitteri et al., 2007), *ERL1* expression is increased in the *er erl2* mutant. In the mutant, we did not observe a statistically significant change in expression of ERf co-receptors, *SERKs*, or of EPFL ligands that do not regulate stomata development (Fig. S2D). EPFL2 might be downregulated by auxin indirectly after prolonged exposure to the hormone (Tameshige et al., 2016), but our results suggest that it is not a direct auxin target.

In response to 30-minute treatment with 1μM IAA, in both wild type and the mutant more than 100 genes were as least 2-fold upregulated and around 20 genes were at least 2-fold downregulated (Fig. 10A and B, Table S2). The main groups of genes upregulated by auxin were Small Auxin Upregulated RNAs (*SAURs*), transcriptional repressors *IAAs*, *GH3* auxin conjugation enzymes, ethylene biosynthesis enzymes *ASCs*, and the Lateral Organ Boundaries (*LBD*) transcription factors. The main group of genes downregulated by auxin were pectin lyases. A comparison of *IAA* and *GH3* gene expression in *er erl2* and the wild type suggested that the mutant can efficiently respond to auxin (Fig.10C and E). The main difference in responses to auxin was the weaker upregulation of *SAURs* in the mutant (Table S2 and Fig. 10D). However, the majority of these *SAURs* are already upregulated in the mutant before the treatment. Auxin treatment equalizes the level of these *SAURs* in the wild type and mutant. We observed that only two auxin upregulated *SAURs*, *SAUR20* and *SAUR34*, were significantly downregulated in the mutant, and while auxin was able to induce them in the mutant, their accumulation never reached the wild type levels. Multiple factors can explain the increased expression of *SAURs*: decreased conjugation of auxin by GH3 auxin-amido synthetases (Fig. 10E), reduced auxin sequestration into the endoplasmic reticulum by auxin efflux carrier PIN5 (Fig.10F), or an increased uptake of auxin from extracellular space by auxin influx carrier LAX1 (Fig. 10F). However, additional experiments are necessary to establish causation.

**Figure 10.**
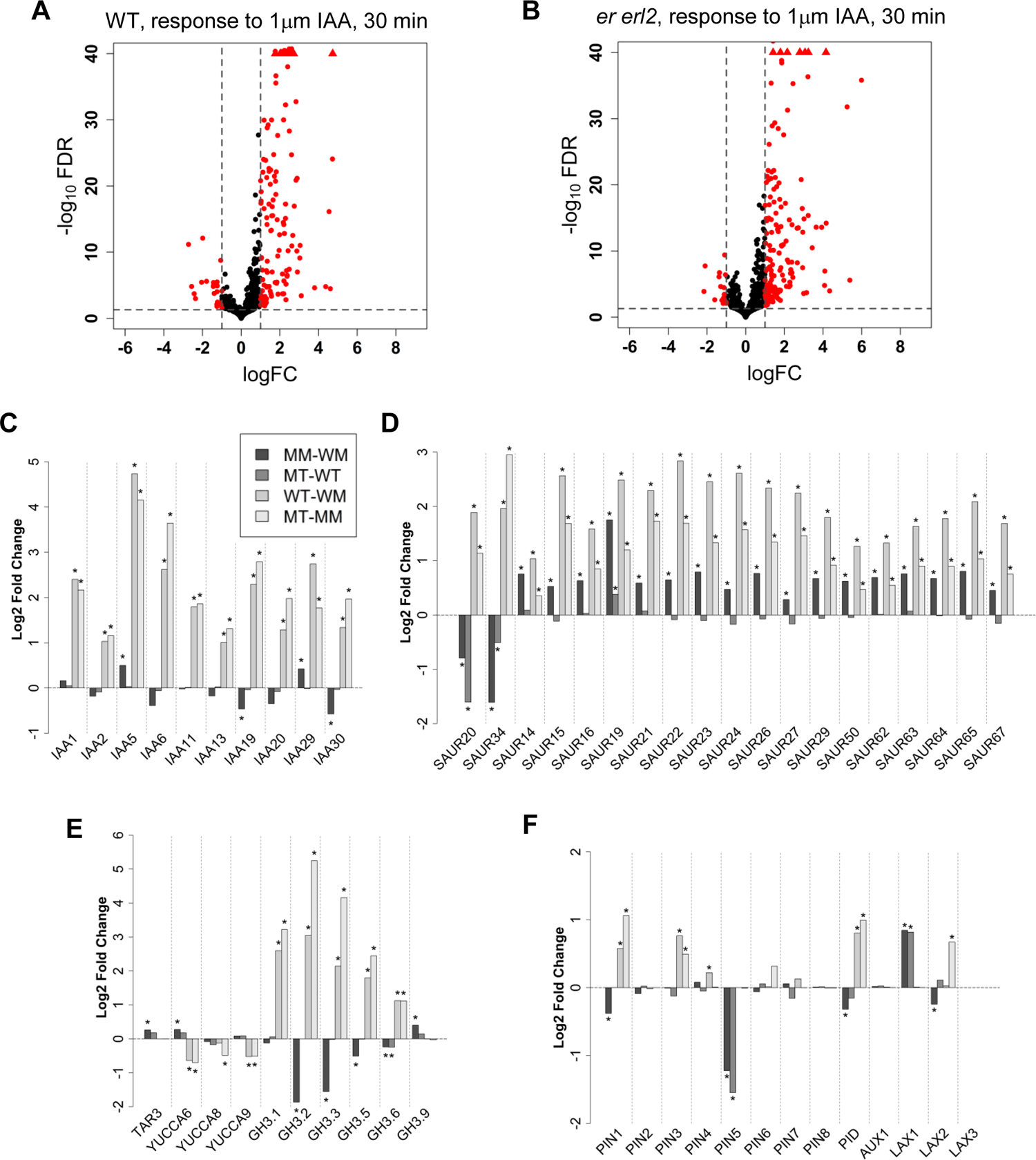
Comparison of direct auxin responses in *er erl2* and the WT. A and B. Volcano plots showing differentially expressed genes (red points) that have a |LogFC| >1 and FDR < 0.05 for **A.** WT-MT and **B.** MT-MM. **C-F** Differential gene expression in all four comparisons (MM-WM, MT-WT, WT-WM, MT-MM) **C.** Auxin responsive IAAs **D.** Auxin responsive SAURs **E.** Auxin biosynthesis and conjugation genes **F.** Auxin transport genes. MM = mutant mock treated, MT = mutant + 30 minutes 1uM IAA treatment, WM = wild type mock treated, WT = wild type + 30 minutes 1uM IAA treatment LogFC = Log 2-fold change. Log 2-fold changes have been shrunk with ashr. * = FDR < 0.05

While it has been proposed that ER promotes auxin biosynthesis (Qu et al., 2017), our results do not support this hypothesis as we observed no or a small opposite effect of ER and ERL2 on auxin biosynthesis enzymes (Fig. 10E). Taken together, our data suggest that ERfs may play some role in fine-tuning auxin metabolism and transport, but they do not play a significant role in the direct transcriptional auxin responses.

## DISCUSSION

### The role of ERf signaling in auxin metabolism

ERf signaling regulates many auxin-controlled processes such as initiation and spacing of leaves and ovules, phototropism, and formation of leaf (Chen et al., 2013; Tameshige et al., 2016; Kawamoto et al., 2020)and the question of how these two signaling pathways are connected has received a fair bit of attention (Chen et al., 2013; Tameshige et al., 2016; Qu et al., 2017; Du et al., 2018; Cai et al., 2021; Liu et al., 2021). Auxin accumulation is increased in the epidermal layer of the *erf* mutants (Chen et al., 2013; Tameshige et al., 2016). However, there is no consensus on the cause and consequences of this increase. Previously, we hypothesized that ERfs regulate leaf initiation by controlling auxin transport (Chen et al., 2013). Other groups proposed that ERfs inhibit auxin responses (Tameshige et al., 2016), promote auxin responses (Cai et al., 2021; Liu et al., 2021), or promote auxin biosynthesis (Qu et al., 2017). The data presented here suggest that all these conclusions, including our own, are at least partially incorrect.

The suggested role of ERfs in regulation of auxin transport is based on the findings that PIN1 and DR5rev:GFP expression are severely decreased in the internal tissues of *er erl1 erl2* (Chen et al., 2013). However, in the SAM PIN1 is essential only in the L1 layer and not in internal tissues, and there mostly for initiation of flowers and not rosette leaves (Guenot et al., 2012; Kierzkowski et al., 2013). Here, we show that in *er erl1 erl2* inhibition of auxin efflux by removing PINs or via an inhibitor of polar auxin transport partially rescues leaf initiation. Therefore, decreased expression of PINs in the internal layers of the mutant cannot be the cause of decreased leaf initiation.

Previous research suggested that ERfs promote expression of *YUCCA* genes and as a result promote auxin biosynthesis (Qu et al., 2017). The epidermis is a tissue layer central to initiation of leaves and to many other auxin responses (Kutschera, 1992; Procko et al., 2016; Kim et al., 2020). A stronger DR5rev:GFP signal in the L1 layer of the *erf* SAM (Chen et al., 2013) and in the epidermis of *erf* leaves (Tameshige et al., 2016) as well as increased expression of *SAUR* genes in *er erl2* point to increased auxin accumulation. In addition, our transcriptomic analysis identified only a subtle increase in expression of two auxin biosynthesis enzymes, TAR3 and YUCCA6, in *er erl2* indicating that ERfs have no or at most a weakly negative impact on auxin biosynthesis. Transcriptomic data suggests several potential causes of altered auxin metabolism. Four *GH3.3* auxin conjugating enzymes are downregulated in *er erl2*. Increased expression of the auxin influx transporter *LAX1* and decreased expression of *PIN5*, which sequesters auxin in the lumen of ER, might be other factors contributing to increased auxin accumulation in the cytosol of *erf* epidermal cells. The negative impact of ERfs on auxin metabolism could be through enhancement of auxin storage as a conjugate or in the endoplasmic reticulum and by reduced uptake of auxin from the extracellular space but not through its biosynthesis.

Surprisingly, increased accumulation of auxin is not sufficient for efficient initiation of leaves in the *er erl1 erl2* SAM, and an increase in auxin responses stimulated by expression of the modified auxin receptor ccvTIR1 can only partially rescue leaf initiation in the mutant. This fact prompted us to investigate whether ERfs, similar to brassinosteroids and other signaling pathways, regulate a specific auxin signaling output (Cho et al., 2014; Chandler, 2016; Tian et al., 2018). However, transcriptomic analysis suggested that the wild type and *er erl2* mutant respond to auxin similarly. Based on our finding we hypothesize that ERfs promote leaf primordia initiation not through direct impact on auxin biosynthesis, transport, or primary transcriptional responses, but instead by regulation of downstream targets that are also controlled by auxin (Fig. 11).

**Figure 11.**
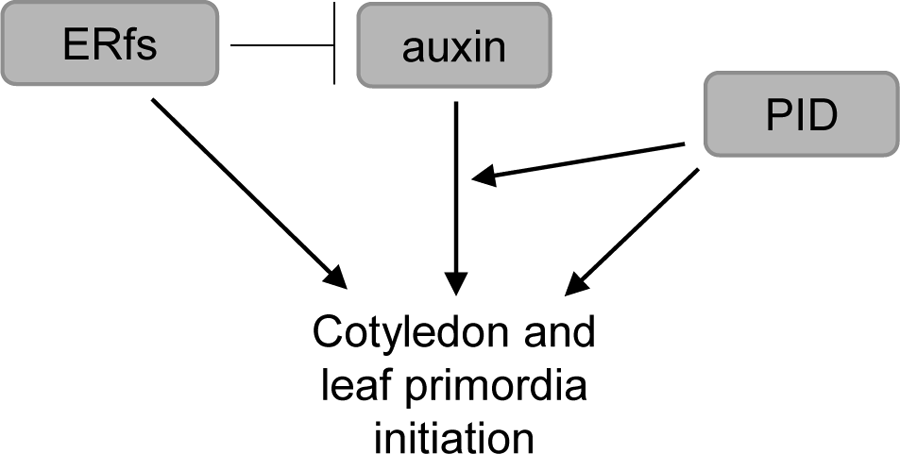
Model of ERfs, auxin and PID function in initiation of aboveground organs. ERfs and auxin promote cotyledon and leaf initiation by targeting a common set of genes. PID promotes initiation of aboveground organs by altering auxin transport and through an auxin independent pathway.

### Initiation of cotyledons and leaf primordia is controlled by multiple signals

Accumulation of auxin in the L1 layer of the SAM is the first sign of leaf primordium initiation. Auxin promotes organogenesis only in a narrow ring of receptive SAM cells where it regulates radial positioning of organs. In a classic experiment, when lanolin paste with IAA was placed on the pin-shaped SAMs, organ primordia always arose at the same distance from the SAM center independent of paste placement (Reinhardt et al., 2000). The amount of auxin in the ring of receptive cells is usually low and transport of auxin into maxima is necessary to reach a threshold concentration. When a uniformly high concentration of auxin was created throughout the ring by placement of IAA paste at the top of the SAM, all receptive cells were induced into organogenesis and a cup-shaped leaf primordium was formed (Reinhardt et al., 2000). Similarly, when we increased auxin signaling evenly throughout the L1 layer of the SAM by expressing the modified auxin receptor ccTIR1, all cells in a narrow ring were induced into organogenesis and a cup-shaped leaf was formed.

What defines the ring of receptive cells in the SAM is still an open question. Multiple findings indicate an increase in auxin perception in the periphery of the SAM (Hardtke and Berleth, 1998; Vernoux et al., 2011; Ma et al., 2019; Galvan-Ampudia et al., 2020). However, an increase of auxin sensing in the center of the meristem does not lead to formation of organs there (Ma et al., 2019). It was hypothesized that an auxin-independent mechanism supports organogenesis in the vegetative meristem (Guenot et al., 2012). One of the mechanisms could be a spatial distribution of abaxial and adaxial genes in the SAM (Caggiano et al., 2017). Our analysis of *er erl1 erl2* and *er erl1 erl2 pid* mutants suggests that ERfs promote leaf initiation in parallel to auxin. Both mutants accumulate and sense auxin in the L1 layer of the SAM, but organogenesis is decreased or absent. Because an increased auxin concentration can partially rescue the *er erl1 erl2* mutant, we hypothesize that ERfs and auxin have common downstream targets in the SAM.

PID is known to promote cotyledon development synergistically with its paralogs WAG1 and WAG2 as well as with several other genes: PIN1, ENP/MAB4, NCP1/AtMOB1, YUC1, and YUC4 (Furutani et al., 2004; Treml et al., 2005; Cheng et al., 2007; Dhonukshe et al., 2010; Cui et al., 2016). It is hypothesized that the synergistic function of these genes is due to their involvement in the formation of auxin maxima. However, this has been shown experimentally only for the *pid ncp1* mutant, where expression of DR5rev:GFP is decreased in the areas of cotyledon formation (Cui et al., 2016). Inexplicably, the *pid enp* mutants often accumulate auxin in the SAM but cannot form cotyledons (Treml et al., 2005). Here we demonstrate that ERf is yet another group of genes that promote cotyledon formation synergistically with PID. Uniquely, ERfs have synergy with PID not only in cotyledon initiation but also in initiation of leaves. The quadruple *er erl1 erl2 pid* mutant accumulates auxin into maxima in the periphery of the SAM, but auxin is unable to induce outgrowth of either cotyledons or leaves. This suggests that other factors in addition to auxin are necessary for cotyledon and leaf initiation. Our data suggest that the role of PID in organ initiation cannot be limited to regulation of auxin transport, and PID is likely to have multiple targets, a characteristic typical for a kinase. It has been proposed that PID interacts with the adaxial HD-ZIP III transcription factor REVOLUTA (Liu et al., 2019).

However, whether these interactions are biologically significant has not yet been determined. In the *pin1 pid* mutant, cotyledon development can be rescued by mutations in either *CUP-SHAPED COTYLEDON* genes (*CUCs*) or *SHOOT MERISTEMLESS* (*STM*) suggesting that CUCs and STM function downstream of PIN1 and PID (Furutani et al., 2004). It would be interesting to investigate in the future whether organogenesis in the *er erl1 erl2 pid* mutant can be rescued by mutations in these genes, and whether ERfs and PID play any role in regulation of CUCs and STM expression.

### Unequal contribution of ERfs and auxin to leaf and flower initiation

It is often assumed that the molecular mechanisms of leaf and flower initiation are identical. While auxin is central to both processes, the finer details of leaf and flower initiation differ. For example, *pin1*, *pid,* and *mp* mutations have a considerably milder impact on leaf initiation compared to flowers (Okada et al., 1991; Bennett et al., 1995; Przemeck et al., 1996; Gälweiler et al., 1998; Barkoulas et al., 2008; Guenot et al., 2012) which suggests that in some species accumulation of auxin into peaks is not a limiting factor during leaf initiation. At the same time, auxin is a limiting factor in the inflorescence meristem, and its increased accumulation in *er erl2* rescues organ initiation in *pin1*. In contrast, ERfs mainly control leaf initiation and have only a moderate impact on flower phyllotaxy (Chen et al., 2013). Knowledge of ERf downstream targets might help us to understand the difference in requirements for initiation of organs during vegetative and reproductive stages.

## Materials and Methods

### Plant Materials and Growth Conditions

The *Arabidopsis thaliana* ecotype Columbia was used as the wild type. *Er erl1 erl2* (Shpak et al., 2004), *pin1-201* (De Rybel et al., 2009), *pin2/eir1-1* (Roman et al., 1995), *pin3-5* (Žádníková et al., 2010), *pin4-3* (Friml et al., 2002), *pin7-2(Friml et al., 2003)*, and *pid-3 (CS8064)* (Bennett et al., 1995) mutants have been described previously. The quadruple *er erl1/+ erl2 pin* mutants were generated by crossing *pin1-201/+, pin2/eir1-1, pin3-5, pin4-3, pin7-2* with *er erl1/+ erl2*. The following primers were used for genotyping: for *pin1-201* – pin1-5 (5’-GTCCCTCATTTCCTTCAAGTACTGTT-3’), pin1-6 (5’-ACCACCACAACATTAACATCATCATTAT-3’), and LBB1 (5’-GCGTGGACCGCTTGCTGCAACT-3’); for *pin3-5* – pin3 (5’-CCCATCCCCAAAAGTAGAGTG-3’), pin3.rc (5’-GGAAGTGTGGAGAGGGAAAAG-3’), and LBB1; for *pin4-3* – pin4.984 (5’-CAACGCCGTTAAATATGG-3’), pin4.1837.rc (5’-TTCCCACTACAATTATTCC-3’), and En91R (5’-TGCAGCAAAACCCACACTTTTACTTC-3’); for *pin7-2* – pin7 (5’-TTTACTTGAACAATGGCCACAC-3’), pin7.rc (5’-GGTAAAGGAAGTGCCTAACGG-3’), and LBB1.3 (5’-ATTTTGCCGATTTCGGAAC-3’).

The *pin2* plants were identified based on their agravitropic roots by growing seedlings vertically for three to five days and then rotating the plate 90^0^. The *erl1-2* and *erl2-1* mutations were genotyped as described previously (Chen et al., 2013). During measurements of leaf initiation, analyzed plants were the progeny of *pin er erl1/+ erl2* and quadruple mutants were identified using the stomata clustering phenotype as a mark of homozygosity for *erl1-2.* In the case of *pin1 er erl1 erl2*, plants were the progeny of *pin1/+ er erl1/+ erl2* and we used altered number or placement of cotyledons as a mark of homozygosity for *pin1*. The *pid er erl1 erl2* mutant was generated by crossing *pid/+* with *er erl1/+ erl2*. The *pid-3* mutant is the result of a point mutation L226F (Christensen et al., 2000) and the genotype of plants was determined by amplifying a segment of PID with RT-PID-c783F (5’-TTATGCCGCCGAAGTTCTAGTG-3’) and PID6 (5’-GACGAGGAAGATTCAACGGCT-3’) primers and then digesting it with the MnlI restriction enzyme. The *pid-3* mutation abolishes the MnlI site.

Plants expressing DR5rev:GFP in the wild type and in *er erl1 erl2* have been described previously (Benková et al., 2003; Chen et al., 2013). The *pin1-201* and *pid-3* plants with DR5rev:GFP were selected in the F2 generation by the presence of fluorescence and altered cotyledon number or shape after a cross between the *pin1-201/+* or *pid-3/+* and wild type plants expressing DR5rev:GFP. To generate the *pid er erl1 erl2* DR5rev:GFP and *pin1 er erl1 erl2* DR5rev:GFP plants, *pid+/- er erl1+/- erl2* and *pin1+/- er erl1+/- erl2* were crossed with *er erl1+/- erl2* DR5rev:GFP.

For all experiments, seeds were stratified for 2 days at 4°C before they were moved to the growth room. On soil, plants were grown as described previously (Kosentka et al., 2017). For analysis of cotyledon number, seedlings were grown on modified solid Murashige and Skoog (MS) medium supplemented with 1% (w/v) sucrose. For analysis of leaf initiation, plants were grown either on solid or liquid modified MS media supplemented with 1% (w/v) sucrose.

Seedlings were grown in liquid media with gentle agitation using a nutating rocker. On average, growth on liquid MS media was slower compared to solid MS media. The following chemicals were added to liquid MS media as indicated: 1, 10 or 100 μM L-Kynurenine (Kyn) (VWR); 0.5, 1, 5, 10 μM N-1-naphthylphthalamic acid (NPA) (Chem Service, Inc. PS-343); 0,5, 1,5,10,100 μM 2,4-Dichlorophenoxyacetic acid (2,4D) (Sigma Aldrich). The stock solutions of L-Kynurenine, NPA and 2,4D were prepared using DMSO. For all mock treatments an equal volume of DMSO was added. The *pid er erl1 erl2* mutants were also germinated and grown on modified MS medium plates supplemented with 10 μM 2,4-D. At 13 DPG, *pid er erl1 erl2* were moved to MS medium plates without 2,4-D and then imaged at 33 DPG.

### Generation and analysis of transgenic plants expressing ccvTIR1

For plant transformation, two plasmids were created with the AtML1 (pDED101) or EPFL4 (pDED103) promoter driving expression of ccvTIR1. At first, the 3xFLAG-ccvTIR1 sequence and the nopaline synthase terminator (NOS) terminator were cut out from pAN19-3xFLAG-ccvTIR1-NOSt plasmid (Addgene plasmid #108546) (Uchida et al., 2018) and inserted into pPZP221 as a BamHI and EcoRI fragment. The created plasmid was named pDED100. The 3.6 kb AtML1 promoter was amplified with the primers AtML1 pr1 (5’-ACATCTCGAGCATTGATTCTGAACTGTACCC-3’) and AtML1pr2.rc (5’-TTAGGATCCAACCGGTGGATTCAGGGAG-3’) using wild type DNA as a template, digested with BamHI and XhoI, and cloned into BamHI/SalI digested pDED100. The 2.8kb EPFL4 promoter was amplified with the primers EPFL4 prXhoI (5’-ATATCTCGAGTAGTTCACCATTTTGGTTG-3’) and Q.EPFL4-2 (5’-AAGAAAGCTGGGTGGATCCTAGTCAAGAACCGGAGAGGA-3’) using previously created plasmid pPZK424 (Kosentka et al., 2019), digested with BamHI and XhoI, and cloned into BamHI/SalI digested pDED100. The pDED101 and pDED103 plasmids were transformed into an *Agrobacterium tumefaciens* strain GV3101/pMP90 by electroporation and introduced into the wild type and *er erl1/+ erl2* plants by the floral dip method. In the T3 generation, we selected lines homozygous for the transgene using gentamicin resistance. The *er erl1 erl2* mutants were identified based on the stomata clustering phenotype. Seedlings were grown on freshly made solid MS media plates with or without 50 μM cvxIAA (Fisher Scientific) for 3 DPG and examined by DIC microscopy.

### Microscopy

Measurements of leaf initiation were performed using DIC microscopy as described previously (Chen et al., 2013). Seedlings were fixed overnight in ethanol: acetic acid 9:1 (v/v), rehydrated with an ethanol series to 30% ethanol (v/v) and then cleared in chloral hydrate solution. The pH of the chloral hydrate (Sigma-Aldrich C8383): water: glycerol 8:1:1 (w/v/ v) solution was adjusted to 4.2 with 10 mM KOH to prevent degradation of young tissues. In our experience, chloral hydrate varies in acidity between different batches.

One-month-old *pid er erl1 erl2* and *pin1 er erl1 erl2* plants were imaged using a Leica MZ6 modular stereomicroscope (Leica Microsystems) with a Leica MC190 HD Microscope Camera (Leica Microsystems). Confocal images were taken using a Leica SP8 White Light Laser Confocal microscope. A 488-nm White Light Supercontinuum Laser (Leica Microsystems) was used to excite the EGFP fluorophore. A ‘UV’ 405 nm diode laser (Leica Microsystems) was used to excite 4’,6-diamino-phenylindole (DAPI) fluorescence. A HyD ‘Hybrid’ Super Sensitivity SP Detector (Leica Microsystems) and a PMT SP Detector (Leica Microsystems) were used to collect emission of EGFP and DAPI, respectively. Z-stacks made with sequential line scanning were used to evaluate 3D expression patterns and to select the 2D image from the SAM center. One cotyledon was removed prior to imaging to expose the SAM of 1 DPG seedlings. Samples were incubated at room temperature in 1 mL of PBS solution containing 1% Triton X-100 and 18 μM DAPI under vacuum for ∼1-2 min and then for ∼7-8 min with gentle rocking without vacuum. Then samples were washed with 1× PBS buffer to remove excess dye for approximately 1 min and imaged.

The Fiji image-processing package was used for all quantitative image measurements. The center slices of 6-8 shoot meristems in mock and NPA treated *er erl1 erl2* seedlings were chosen by analyzing the overall shape of the meristem. Using the Fiji image processing software, the freehand tool was used to draw a line across the L1 cell layer of the SAM starting from one adaxial leaf junction and finishing at the opposite adaxial leaf junction. A plot profile measuring the mean pixel intensity was generated for each seedling measured. The data points were binned into 8 μm increments where 0 μm marks the center of the SAM.

### Transcriptomic Analysis

Three-day-old post germination WT (Col-0) or mutant (*er-105 erl2-1)* seedlings grown on MS solid medium were acclimated to MS liquid media for 30 min and then treated for 30 minutes with either mock (0.02% DMSO) or 1μM IAA (0.02% DMSO). Total RNA for library construction was extracted using a Spectrum Plant Total RNA Kit (Sigma). RNA quality was assessed using an Agilent 4200 TapeStation. Total RNA was poly-A selected and stranded libraries were constructed using a NEBNext Ultra II Directional RNA Library Prep Kit for Illumina. Paired-end 150bp sequencing was performed at Novogene on a Novaseq 6000. Three biological replicates were run for each sample and each replicate was sequenced at a depth of 30M fragments. Read quality control was performed with FastQC 0.11.9. Reads were mapped to the TAIR 10.1 (Lamesch et al., 2012) genome and the Araport 11 annotation (Cheng et al., 2017) using STAR 2.7.6a (Dobin et al., 2013). RSeqc 2.6.4 (Wang et al., 2012) was run on the mapped reads as quality control. Reads were assigned to features using featureCounts 2.0.0 (Liao et al., 2014) in paired-end mode. The subsequent counts matrix was then imported into R 3.6.3 (R Core Team) and differential gene expression carried out by DESeq2 1.26.0 (Love et al., 2014) using a two-factorial design. P-values were adjusted for multiple comparisons by the Benjamini-Hochberg correction (*q* < 0.05). Log fold change shrinkage was performed using ashr 2.2 (Stephens, 2017). GO enrichment analysis was performed using a custom wrapper around topGO 2.38.1 (Alexa and Rahnenfuhrer, 2016).

## Acknowledgments

We thank Jiri Friml for sharing with us seeds of *pin1-201, pin2/eir1-1, pin3-5, pin4-3,* and *pin7-2* mutants. We thank Ming-Kun Chen and Kate Checuga for technical assistance.

## Accession numbers

Arabidopsis Genome Initiative numbers for the genes discussed are as follows: ATML1 (At4g21750), ER (At2g26330), ERL1 (At5g62230), ERL2 (At5g07180), EPFL4 (At4g14723), PID (At2g34650), PIN1 (At1g73590), PIN2 (At5g57090), PIN3 (At1g70940), PIN4 (At2g01420), PIN5 (At5t16530) PIN7 (At1g23080), STOMAGEN (At4g12970). Sequencing data were deposited at NCBI under GEO accession number GSE197154.

### This article contains supporting information

Figure S1. PIN1 and ERfs synergistically promote inflorescence growth.

Figure S2. Comparison of gene expression in *er erl2* and the wild type.

Table S1. The *er erl1 erl2* mutation reduces number of cotyledons in the *pin1* background

Table S2. Transcriptomics data

